# Reconstructing the origins of the space-number association: spatial and number-magnitude codes must be used jointly to elicit spatially organised mental number lines

**DOI:** 10.1101/446930

**Authors:** Mario Pinto, Michele Pellegrino, Fabio Marson, Stefano Lasaponara, Fabrizio Doricchi

## Abstract

In a series of recent studies we have pointed out that the use of contrasting left/right spatial codes, whether indirectly related to number magnitudes through response selection or directly associated to the same magnitudes to guide their spatial positioning on a mental number line, is crucial in eliciting space-number associations (Aiello, 2012; Fattorini et al., 2015; 2016; Pinto et al., 2018). Nonetheless, this conclusion is based on experiments in which spatial and number-magnitudes codes are used jointly during task performance. Here, in a series of unimanual Go/No-Go tasks with intermixed central numerical and pictorial targets, i.e. arrows pointing to the left or to the right, we explore whether spatial codes used in isolation inherently evoke the left-to-right representation of number magnitudes and, vice-versa, whether number-magnitude codes used in isolation inherently evoke the conceptual activation of left/right spatial codes. In a first series of experiments participants were asked to provide unimanual Go/N-Go responses based on instructions that activated only magnitude codes, e.g. “push only if the number is lower than 5 and whenever an arrow appears”, or only spatial codes, e.g. “push only when an arrow points to the left and whenever a number appears”. In a second series of experiments, the same numerical instructions were combined with the request of responding only to arrows in a specific colour, e.g. “push when the number is lower than 5 and whenever a blue arrow appears”. At variance with a recent experiment by Shaki and Fischer (2018), in our experiments no constant association was present between a specific arrow colour and a specific arrow direction. The results of these experiments highlight no space-number congruency effects: e.g. no faster RTs to arrows pointing to the left rather than to the right when participants attend to numbers lower than 5 and, vice-versa, no faster RTs to numbers lower than 5 rather than higher, when participants attend to arrows pointing to the left. Based on these findings it must be concluded that neither space codes used in isolation can elicit a spatial representation of number magnitudes nor number-magnitude codes used in isolation can trigger the activation of spatial codes. Thus, spatial and numerical codes must be used jointly to evoke spatially organised mental number lines.

## 1. Introduction

One of the issues of major interest in numerical cognition is the relationship between the brain representation of number magnitudes and that of space. The most reliable evidence of a functional interaction between numerical and spatial processing is the *Spatial Numerical Association of Response Codes* (SNARC) effect (Dehaene et al. 1993). This effect shows that when healthy participants belonging to left-to-right reading cultures are asked to make judgments on the magnitude of a number (e.g. higher or lower than 5?, Dehaene, et al., 1990) or on its parity (e.g. odd or even?, Dehaene et al., 1993), reaction times (RTs) to small numbers are faster with response keys placed in the left side of space and those to large numbers with keys in the right side.

The SNARC was initially interpreted as deriving from the congruency between the position of response keys and the position that numbers would inherently occupy on a mental number line that is spatially organised in accord with reading habits, that is with smaller numbers positioned to the left of larger ones in left-to-right reading cultures and vice versa in right-to-left ones (Dehaene et al., 2003; Hubbard et al., 2005; Nuerk et al., 2005). This interpretation considers spatial position an intrinsic part of the semantic representation of number magnitudes. Two relatively recent findings seemed to provide support for this interpretation. The first is the deviation toward larger numbers, i.e. toward numbers that are putatively placed on the right side of the MNL, that was initially described in a group of six right brain damaged patients (RBD) with attentional neglect for the left side of space during the mental bisection of number intervals (Zorzi et al., 2002). The second finding is the RTs advantage that healthy participant displayed in the detection of visual targets presented in the in left side of space when these were preceded by small-magnitude Arabic numbers presented at central fixation, and for targets in the right side when these were preceded by large-magnitude numbers, just as if small numbers triggered automatic shifts of attention toward the left side and large numbers toward the right side of space (Attentional SNARC; Fischer et al., 2003). Nonetheless, both these lines of evidence were recently put into question. First, several investigations in RBD patients showed no relationship between the presence and severity of neglect and a bias toward large numbers during the bisection of number intervals (Doricchi, et al., 2005; Doricchi et al., 2009; Loetscher & Brugger, 2009; Loetscher et al., 2010; Rossetti et al., 2011; van Dijck et al., 2011; Pia et al., 2012; Aiello et al., 2012; van Dijck et al., 2012; Aiello et al., 2013). Most important, two independent studies showed that the bias toward large numbers during the mental bisection of number intervals is correlated, independently of spatial neglect, to a numerically equivalent bias toward large numbers placed in the left side of a visual or mental clock-face during the mental bisection of time intervals (Rossetti et al., 2011; Aiello et al., 2012). This finding suggests a deficit in the representation of small numerical magnitudes that is independent from their spatial positioning in the left or the right side of a mental image. Second, several recent investigations run in adequately large samples of participants failed to replicate the original Attentional-SNARC effect (Zanolie & Pecher 2014; van Dijck et al., 2014; Fattorini et al., 2015; Fattorini et al., 2016; Pinto et al., 2018) and demonstrated no relationship between the strength of this effect and the strength of conventional Magnitude or Parity SNARC effects (Fattorini et al., 2015).

Other interpretations of the SNARC effect were also proposed (for review see Cohen Kadosh et al., 2008; Wood et al., 2008; Fattorini et al., 2016). A series of study (Proctor and Cho, 2006; Santens & Gevers, 2008; Gevers et al., 2010) have argued that culturally acquired associations between the concepts “left/right” and “small/large” determine the SNARC. In particular according to Proctor and Cho (2006) a positive polarity is culturally assigned to concepts like “right” and “large,” while a negative polarity is shared by concepts like “left” and “small.” Consequently, it is easier and faster to associate the concept “smaller than 5” with a “left” key response and the concept “larger than 5” with a right key response than creating the opposite set of associations. A series of important experiments by Gevers et al. (2010) provided support to the role of verbal coding by showing that when spatially incongruous verbal labels are assigned to left and right response keys, so that the key on the left side is labelled “right” and that on the right side is labelled “left”, the SNARC effect remains anchored to verbal labels rather than to the actual position of response keys.

A third interpretation sees the SNARC as depending on the use of spatial response codes: the mental left-to-right organization of number magnitudes would in this case be induced by the contrasting “left vs. right” spatial codes that must be used in the selection of motor response associated with number magnitudes (Keus & Schwarz, 2005; Ishihara et al., 2006; Müller & Schwarz, 2007; Fattorini et al., 2015). In this sense, the left-to-right representation of numbers would be a temporary and “ad hoc” adjustment of number representation to the left-to-right right coding of motor responses required by the SNARC tasks (Fattorini et al., 2015; 2016). The setting of this temporary match would be facilitated by the congruency between the spatial codes imposed by the task and the culturally acquired directional-spatial mapping of alphabetical and numerical material linked to reading habits. This response related account has found support in the results of investigations run with Event Related Potentials (ERPs). These have coherently demonstrated that the SNARC arises at a response-selection stage rather than at an earlier perceptual processing stage of number cues (Keus et al., 2005; Gevers, et al. 2006).

In a relatively recent paper (Aiello et al., 2012), we have highlighted that an important source of evidence for interpreting the SNARC and, more generally, the space-number association as depending on the use of left/spatial codes during task performance, comes from studies that have tested the performance of neglect patients in the SNARC task. These studies have homogenously highlighted slower RTs for number magnitudes that are immediately lower than the numerical reference (e.g. 4 with respect to 5) as compared with numbers that are immediately higher than the same reference (e.g. 6 with respect to 5; Vuilleumier et al., 2004; van Dijck et al., 2012; Zorzi et al., 2012). These findings show that when left/right response codes must be associated with numbers, a truly spatial representation of number magnitudes is elicited and the ipsilesional attentional bias suffered by neglect patients is put in evidence, so that the processing of numbers smaller than the reference is delayed, just as the processing of visual stimuli presented in the neglected left side of space. It is important to note that the same does not happen in the mental bisection of number intervals where no use of left/right response code is required and the numerical bias is no more linked to the presence or severity of spatial neglect (Doricchi, et al., 2005; Doricchi et al., 2009; Loetscher & Brugger, 2009; Loetscher et al., 2010; Rossetti et al., 2011; van Dijck et al., 2011; Pia et al., 2012; Aiello et al., 2012; van Dijck et al., 2012; Aiello et al., 2013). Based on this evidence, we have originally proposed (Aiello et al., 20012) and confirmed in a series of behavioural and EEG studies (Fattorini et al., 2015; 2016; Pinto et al., 2018), that the space-number association does not rely on inherent semantic properties of number magnitudes but, rather, on the use of spatial codes in the numerical task at hand.

In line with our proposal, a number of recent investigations suggest that SNARC-like effects can also be produced when contrasting left/right codes are conceptually associated with number stimuli in a direct manner, rather than indirectly through the selection of motor responses. In a series of studies (Fattorini et al., 2015; 2016; Pinto et al., 2018) we have demonstrated that when left/right codes are used to guide the mental positioning of numerical cues to the sides of a central numerical reference, “5”, the detection of visual targets presented at spatial positions corresponding to that occupied by numerical cues in mental space is facilitated (Spatial-Attentional SNARC). At the electrophysiological level, the active mental positioning of numbers to the left or the right of the central reference is a reflected in an enhancement of facilitatory-preparatory brain activity over the hemisphere contralateral to the mental number position, i.e. Lateral Directing Attention Positivity (LDAP), and in the enhancement of early EEG responses, C1 component, to targets that are presented on the side of visual space that is congruent with that of numerical cues in mental space. No comparable effects are present when numerical cues are passively perceived or when they are classified as function of their magnitude, i.e. lower or higher than “5”. These findings provide support to the hypothesis first advanced in Aiello et al. (2012) and re-proposed in Fattorini et al. (2015; 2016) that the representation of number magnitudes has no inherent left-to-right spatial organization and that this organization is rather temporarily elicited by the use of left/right spatial codes, whether linked to number stimuli through response selection or directly conceptually associated to the same stimuli in the task at hand. In line with our original proposal, Fischer and Shaki (2017) have recently showed that when left/right spatial codes are associated with magnitude codes in the instructions that regulate the release of unimanual Go responses to intermixed numerical targets and pictorial shape-targets pointing to the left or to the right, reaction times (RTs) are faster when in task instructions spatial codes are congruent with the position that numerical targets would occupy on a horizontally oriented MNL. Therefore, an instruction like “*push when the number is lower than 5 or the shape points to the left*” produces faster RTs than an instruction like “*push when the number is lower than 5 or the shape points to the right*”. These results show that the same conceptual associations of spatial and magnitude codes that determine RTs advantages in response selection during the SNARC task, i.e. left-lower than 5 and right-larger than 5, can also produce RTs advantages during a unimanual Go/No-GO stimulus classification task that requires no contrasting spatial codes for response selection. Though interesting, these findings (that, by the way, we were able to replicate, see Supplementary Experiment 1 in Supplementary Material) are based on a task that, like in Fattorini et al. (2015; 2016) and Pinto et al. (2018), still requires the joint use of spatial and magnitude codes. Therefore, these observations still skip one more fundamental question in the search of the origins of the space-number association: do contrasting spatial codes used in isolation inherently evoke the conceptual left-to-right representation of number magnitudes and, vice-versa, do contrasting number-magnitude codes used in isolation inherently evoke the conceptual activation of contrasting left/right spatial codes?

In the present study, we have addressed these questions. In a first experiment with a Go/No-Go task (Experiment 1), in different trials low- or high-magnitude Arabic numbers were alternated at central fixation with left- or right-pointing directional-arrows. In different experimental conditions, participants were asked to: a) provide unimanual Go speeded detection responses only to low- or to high magnitude numbers while responding, with the same response key, to all arrows independently of their direction; b) provide unimanual Go speeded detection responses only to left- or to right-pointing arrows while responding, with the same response key, to all numbers independently of their magnitude. Therefore, in both experimental conditions the task required an intra-categorical judgement for one stimulus category (numbers or letters) and the simple detection of targets belonging to the remaining category. If the isolated contrast between spatial codes elicits an automatic spatial representation of number magnitudes and/or the isolated contrast between number-magnitude codes automatically triggers the activation of spatial codes, then space-number congruency effects should be observed in all experimental conditions. In contrast, if this congruency effects depend on the combined, rather than isolated, use of spatial and number codes, no congruency effects should be found in the same experimental conditions.

In a second series of additional control experiments (Experiments 2 and 3) we have also addressed the results of a recent experiment by Shaki and Fischer (2018) that, contrary to previous studies (Zanolie and Pecher, 2014; Fattorini et al., 2015: Fattorini et al., 2016; Pinto et al., 2018), seem to support the idea that numerical-magnitude codes used in isolation can trigger the activation of spatial codes and the generation of spatially organised mental number lines. In that experiment (Experiment 2 in Shaky and Fischer, 2018), in different trials central numerical targets of different magnitude, i.e. 1 to 9 excluding 5, were alternated with central coloured arrows that pointed left, right, up or down. Participants had to provide unimanual Go/No-Go responses basing on combined number-colour instructions, e.g. push when the number is lower than 5 and when the arrow is red. Shaki and Fischer (2018) found that attending to magnitudes lower than 5 speeded-up RTs to arrows pointing to the left while attending to numbers higher than five speeded-up RTs to arrows pointing to the right, even if arrow direction was irrelevant to the task. Based on these congruency effects between number magnitude and arrow direction, these authors concluded that magnitude codes inherently activate corresponding spatial-codes, i.e. left/right or up/down, thus generating the space-number association, even when, as in the case of their task, the attention of participants is focused on arrow-colour rather than on arrow-direction. In principle, this conclusion could be considered correct on condition that in the task no association is present between a specific arrow-colour and a specific arrow-direction. This is easily obtained when, for example, a red-arrow points to the left or to the right on an equivalent number of trials. Unfortunately, in their experimental task Shaki and Fischer (2018) associated a specific arrow-colour with a specific arrow-direction) so that, for example, all arrows pointing to the left were always and only red while those pointing to the right were always and only green (see section 3.3 in Shaky and Fischer, 2018: “*Arrow colors were fixed for each participant (e.g. red left arrows and green right arrows)*”. It is easy to argue that in this condition an healthy observer that must provide a Go/No-Go response only to red arrows quickly learns that “red” is equivalent to “left”, so that the instruction “push when the number is lower than 5 and when the arrow is red” becomes also equivalent to “push when the number is lower than 5 and the arrow points left” thus re-introducing the use of spatial codes in the task. As a consequence, this experimental condition guarantees in no way the selective activation of magnitude codes, and precludes testing properly whether these codes used in isolation can trigger the activation of directional left/right spatial codes. To better re-address this problem, we run a second series of experiments (2 and 3) with Go/No-Go tasks in which arrow-stimuli pointing to the left or to the right were both depicted with equal probability in one of the two target colours, i.e. blue or yellow, so that no specific association was present between an arrow-colour and an arrow-direction. In a first experiment (Experiment 2), the direction of arrows was randomly alternated from left to right over different trials. In a second experiment (Experiment 3) the left or the right direction of arrows was kept constant in each block of trials. This latter experimental manipulation served to verify whether the spatial code that was constantly and implicitly conveyed by arrow-targets during each block of trials was eventually elaborated and integrated in the mental task set, so to induce space-number congruency effects.

## 2. General Method

### 2.1 Apparatus

All experiments were run in a sound attenuated room with dim illumination. Stimuli were presented on a 15-inch-color VGA monitor. An IBM-compatible PC running MATLAB software controlled the presentation of stimuli and the recording of responses. Participants had their head positioned on a chin rest at a viewing distance of 57.7 cm from the screen. All participants had normal or corrected to normal vision and were naive to the experimental hypothesis of the experiment. Different samples of participants were included in the four experiments of the present study.

### 2.2 Assessment of handedness and counting direction style

All participants were right-handed and at the end of experimental sessions were administered with two tasks that tested their counting direction preference. In a first task (Fischer and Shaki, 2017), four identical black cardboard circles with a diameter of 4 cm were presented equally spaced in a linear array on a blank A4 piece of paper that was positioned in landscape format with its centre aligned to the head-body midsagittal plane of participants. Each participant was asked to “count these circles aloud and touch each circle while counting.” No demonstration was given and the participant’s order of counting was recorded in a single trial by the experimenter as being directed either from left to right or vice-versa. In a second task (Lindemann, et al., 2011) each participant was asked to hold hands in front of her/his body/head and count aloud from one to ten, using fingers to count. Similarly to the former task, the examiner scored whether the count proceeded from the left to the right hand or vice-versa. In all experiments, we run a series of analyses to investigate the possible influence of counting direction style on space-number congruency effects. To this aim in each experiment, we compared RTs performance between the two subgroups of participants who had left-to-right vs. right-to-left counting direction style.

## 3. Experiment 1

### 3.1 Participants

Twenty-four healthy right-handed student (16 females, 8 males; mean age = 22,8 years, SD = 2,9 years) from the University “La Sapienza” in Rome participated in the experiment. All participants showed left-to-right scanning in the inspection of linear arrays of four cardboard circles (Fischer and Shaki, 2017). On finger counting test (Lindemann et al., 2011) 11 participants showed left-to-right and 13 right-to-left preference.

### 3.2 Procedure

Each trial started with the 500 ms presentation of a central fixation cross (1.5° × 1.5°). At the end of this delay an Arabic digit (1, 2, 8 or 9; size = 1.5° × 1°; font = Arial) or an arrow coloured in blue or yellow and pointing to left or to the right (size = 1.5° × 0.8°), replaced the central fixation cross. There was no specific association between arrow-colour and direction, so that over the total number of trials both yellow and blue arrows pointed an equivalent number of times to the left or to the right. Target stimuli remained available for response for 2000 ms. The inter-trial interval was 500 ms. This timing of trial events is equivalent to that used in Shaki and Fischer (2018). To investigate whether spatial codes used in isolation implicitly generate the left-to-right representation of number magnitudes or, vice-versa, whether number-magnitude codes used in isolation implicitly trigger the conceptual activation of left/right spatial codes, each participant used the following four rules, that corresponded to four different experimental conditions, to provide Go/No-Go responses to numerical- or arrow-targets: a) press the spacebar if an arrow points to the left and whenever a number appears; b) press the spacebar if an arrow points to the right and whenever a number appears; c) press the spacebar if the number is smaller than 5 and whenever an arrow appears; d) press the spacebar if the number is larger than 5 and whenever an arrow appears.

The four experimental conditions were administered during a single experimental session. The order of experimental conditions was counterbalanced among participants. The task was divided into 4 blocks of trials, each corresponding to a different experimental condition. Each block consisted of 256 trials, 128 with numerical-targets (i.e. 32 trials per Arabic number) and 128 with arrow-targets (32 trials per each of the four colour-direction combination). A short break was allowed between blocks. At beginning of the experimental session, participants performed a training session composed of 16 trials.

### 3.4 Results

Go trials in which no response was provided (misses) or in which RTs were above 1000 ms or below 100 ms were not included in the analyses. This procedure was applied to all the analyses summarized in the present report.

4,9 % of trials were discarded from the analyses. RTs to numerical targets, obtained in the Experimental Session “a” and “b”, were entered in an Experimental Condition (Congruent, Incongruent) × Target Magnitude (Smaller, Larger) ANOVA. The Experimental Condition effect was not significant [F (1, 23) = 2.83, p = .11, η_p_^2^ = .12], thus showing no number-space association (SNAs). In addition, the Experimental Condition × Target Magnitude was not significant [F (1, 23) = 1.91, p = .18, η_p_^2^ = .08; Figure 1A], The presence of SNAs was also investigated through regression analysis (Lorch& Myers, 1990). We first estimated individual differential RTs (dRTs) obtained by subtracting RTs to Small magnitude and Large magnitude numerical targets produced when participants attended to Left pointing arrows from equivalent RTs produced when attending to Right pointing arrows. Digit magnitude was used as predictor variable and dRTs as criterion variable. With this method, a significant negative slope indexes the presence of the SNAs (Fias et al., 1996; Ito &Hatta, 2004). One-sample t-tests showed that the Slope describing the linear regression did not differ from zero [t (23) = −1,67, p = .11; average = −1,16; SD = 3.28; Figure 1B]: this result confirms the absence of the SNAs. The counting direction style Group × Experimental Condition interaction was far from being significant [F (1, 22) = .39, p = .54, η_p_^2^ = .01]: this show no difference in the lack of space-number congruency effects between the subgroup of participants that counted from left to right (n = 11) and those who counted from right to left (n = 13).

**Figure 1.**
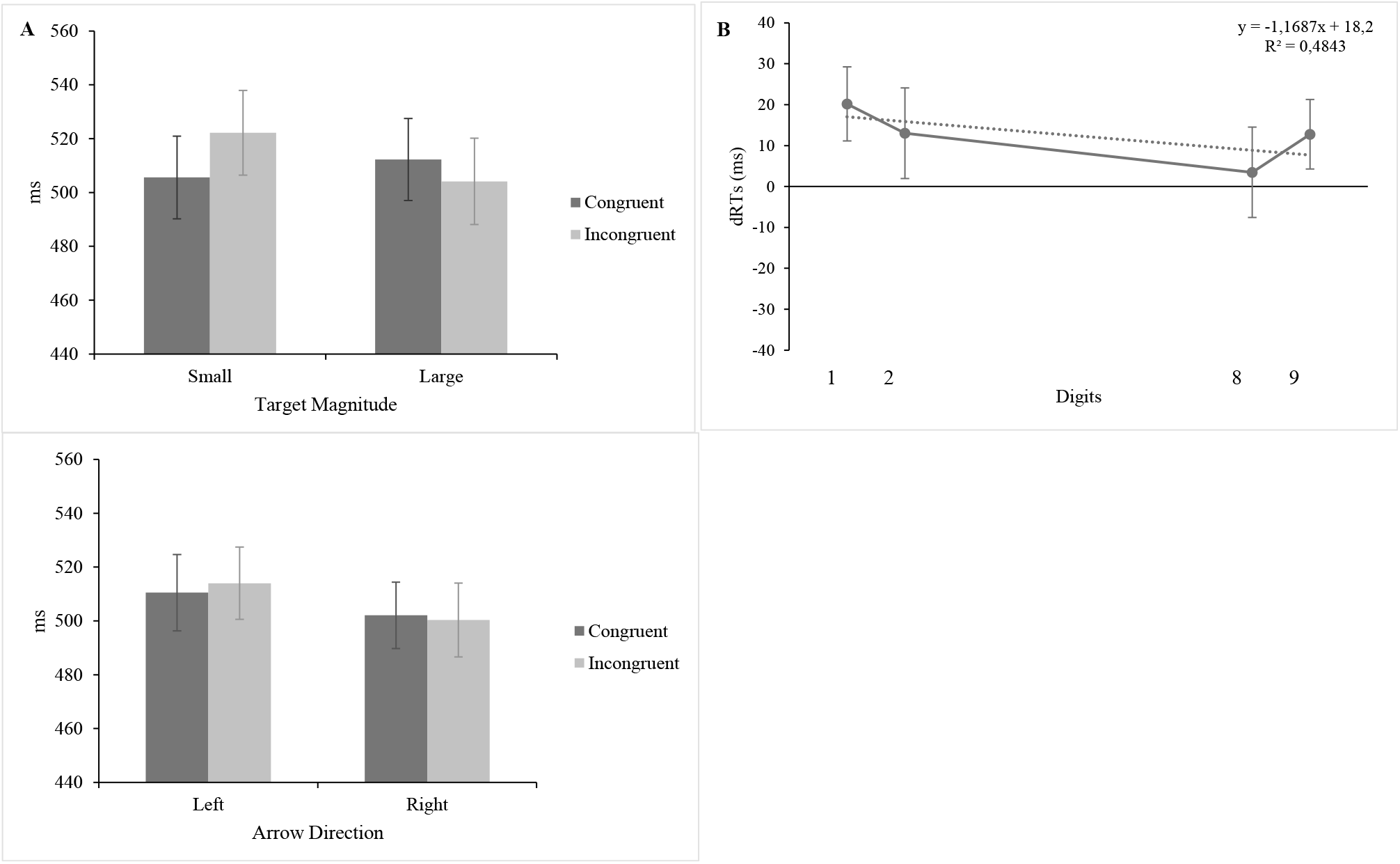
*Experiment 1:* (**A**) Average RTs (with SE) to Small magnitude (1,2) and Large magnitude (8,9) numerical targets in the Congruent task condition (RTs to Small number magnitudes when participants must respond only to Left pointing arrow targets and RTs to Large number magnitude when participants must respond only to Right pointing arrows) and in the Incongruent condition (RTs to Small number magnitudes when participants must respond only to Right pointing arrow targets and RTs to Large number magnitude when participants must respond only to Left pointing arrows). (**B**) Regression slope describing differential RTs (dRT in ms) to Small magnitude and Large magnitude numerical targets while attending to Right pointing minus Left pointing arrows. (**C**) Average RTs (with SE) to Left and Right pointing arrows in the Congruent task condition (RTs to Left arrows when participants must respond only to Small number magnitudes and RTs Right pointing arrows when participants must respond only to Large number magnitudes) and in the Incongruent condition (RTs to Left arrows when participants must respond only to Large number magnitudes and RTs to Right pointing arrows when participants must respond only to Small number magnitudes).

Finally, we entered RTs to arrows targets, obtained in the Experimental Session “c” and “d”, in an Experimental Condition (Congruent, Incongruent) ×Arrow Direction (Left, Right) × Arrow Colour (Yellow, Blue) ANOVA, The Experimental Condition effect was not significant [F (1, 23) = .05, p = .82, η_p_^2^ < .01; Congruent: 506 ms, Incongruent: 507ms; Figure 1C]. This results shows that RTs to Left arrows when participants must respond only to Small number magnitudes and RTs Right pointing arrows when participants must respond only to Large number magnitudes (Congruent condition), are not faster from RTs produced in response to Left arrows when participants must respond only to Large number magnitudes and RTs to Right arrows when participants must respond only to Small number magnitudes. The Arrow Colour had no influence on RTs [F (1, 23) = 2.36, p = .15, η_p_^2^= .09; Yellow Arrows: 505 ms; Blue Arrows 508 ms].

Also in this case no difference was found as function of counting direction style [Group × Experimental Condition interaction: F (1, 22) = .72, p= .40, η_p_^2^= .03].

## 4. Experiment 2

### 4.1 Participants

Twenty healthy right-handed student (10 females, 10 males; mean age = 23,7 years, SD = 1,9 years) from the University “La Sapienza” in Rome participated in the experiment. All participants showed left-to-right scanning in the inspection of linear arrays of four cardboard circles (Fischer and Shaki, 2017). On finger counting test (Lindemann et al., 2011) 10 participants showed left-to-right and 10 right-to-left preference.

### 4.2 Procedure

Each trial started with the 500 ms presentation of a central fixation cross (1.5° × 1.5°). At the end of this delay an Arabic digit (1, 2, 3, 4, 6, 7, 8, 9; size = 1.5° × 1°; font = Arial) or a coloured arrow (blue or yellow, pointing to left or right; size = 1.5° × 0.8°), replaced the central fixation cross. Stimuli remained available for response for 2000 ms. The inter-trial interval was 500 ms.

As in experiment 1, and contrary to Shaki and Fischer (Exp. 2, 2018), there was no specific association between arrow-colour and arrow-direction. At variance with experiment 1, where each experimental condition required only the discrimination of number-magnitude or arrow-direction, in each of the four different experimental conditions/blocks of trials of experiment 2, participants had to discriminate both the magnitude of numerical targets and the colour of arrow-targets according to one of the following instructions: a) press the spacebar when the number is smaller than 5 and when the arrow is yellow; b) press the spacebar when the number is smaller than 5 and when the arrow is blue; c) press the spacebar when the number is larger than 5 and when the arrow is yellow; d) press the spacebar when the number is larger than 5 and when the arrow is blue.

The four experimental conditions were administered during a single experimental session. The task was divided into four different blocks of trials, one for each experimental condition. The order of experimental conditions/blocks was counterbalanced among participants. Each block consisted of 256 trials, 128 with numerical-targets (i.e. 16 trials for each Arabic number) and 128 with arrow-targets (32 trials for each of the four colour-direction combination). A short break was allowed between blocks. At beginning of the experimental session, participants performed a training session composed of 16 trials.

### 4.3 Results

1,8 % of trials were discarded from the analyses. We entered RTs to arrows targets in an Experimental Condition (Congruent, Incongruent) × Arrow Direction (Left, Right) × Arrow Colour (Yellow, Blue) ANOVA. The Experimental Condition effect was not significant [F (1, 19) = .80, p = .38, η_p_^2^ = .05: Congruent: 485 ms; Incongruent: 471 ms, Figure 2]: this shows that attending to numbers lower than 5 did not speed up RTs to arrows pointing left and, vice versa attending to numbers higher than 5 did not speed up RTs to arrows pointing right. Arrows Colour had no influence on RTs [F (1, 23) = .03, p = .87, η_p_^2^<.01; Yellow Arrows: 485 ms; Blue Arrows 487 ms].

**Figure 2.**
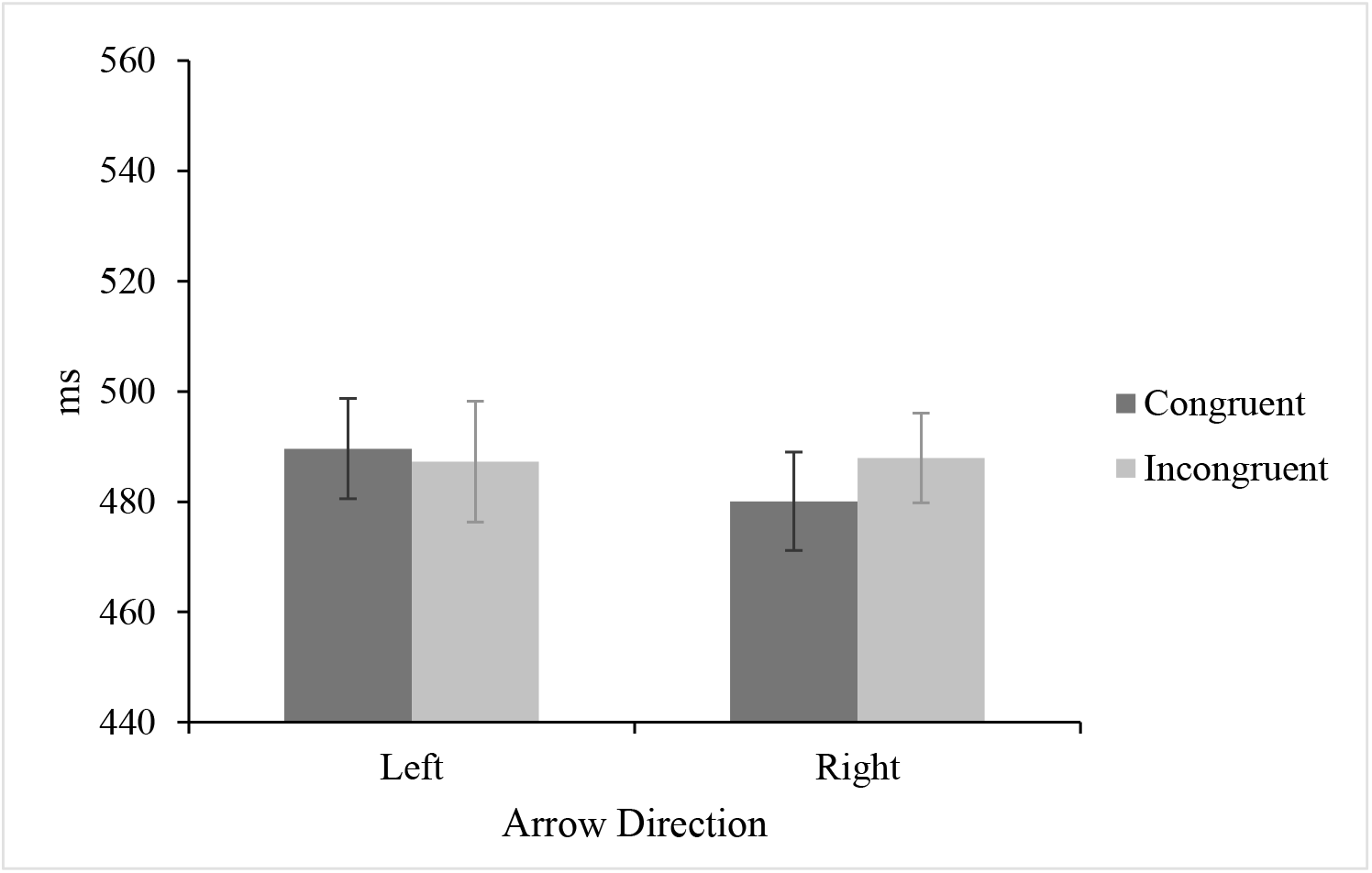
*Experiment 2:* In this experiment, in different blocks of trials participants provided Go responses only to blue or yellow arrow targets. No processing of arrow-direction was required by the task. Blue and yellow arrow-targets pointed with equal probability in the Left or Right direction. Average RTs (with SE) to Left and Right pointing arrows in the Congruent task condition (RTs to Left arrows when participants must respond only to Small number magnitudes and RTs Right pointing arrows when participants must respond only to Large number magnitudes) and in the Incongruent condition (RTs to Left arrows when participants must respond only to Large number magnitudes and RTs Right pointing arrows when participants must respond only to Small number magnitudes).

The counting style direction Group × Experimental Condition interaction [F (1, 18) = 1.34, p= .27, η_p_^2^ = .08]: highlighted no difference in space-number congruency effect between participants counting left to right (n = 10) and those counting from right to left (n = 10).

## 5. Experiment 3

### 5.1 Participants

Twenty healthy right-handed student (10 females, 10 males; mean age = 22,9 years, SD = 1,6 years) from the University “La Sapienza” in Rome participated in the experiment. All participants showed left-to-right scanning in the inspection of linear arrays of four cardboard circles (Fischer and Shaki, 2017). On finger counting test (Lindemann et al., 2011) 10 participants showed left-to-right and 10 right-to-left preference.

### 5.2 Procedure

Each trial started with the 500 ms presentation of a central fixation cross (1.5° × 1.5°). At the end of this delay an Arabic digit (1, 2, 8, 9; size = 1.5° × 1°; font = Arial) or an colour arrow (blue or yellow pointing to left or right; size = 1.5° × 0.8°), replaced the central fixation cross. Stimuli remained available for response for 2000 ms. The inter-trial interval was 500 ms.

In experiment 2 we demonstrated that, like in experiment 1, when in the task there is no systematic association between arrow-colour and arrow-direction and an equal number of left- and right-pointing arrow targets are presented, the processing of a specific numerical magnitude does not speeds up the detection of arrows that point in the direction of the putative position occupied by the same numerical magnitude on a horizontally arranged MNL. This result shows that the use of magnitude codes does not inherently trigger the activation of corresponding left/right spatial codes. In experiment 3, using the same instructions/experimental conditions of experiment 2 we wished to test whether the activation of spatial codes is facilitated when the lateral direction of arrows is kept constant within the same block of trials. To this aim, in the task used in the Experiment 3 half of participants were asked to respond only to yellow arrows according to the each one on the following four types of instructions/experimental conditions: a) “press the spacebar when the number is smaller than 5 and when the arrow is yellow”: only left-pointing arrows are presented; b) “press the spacebar when the number is larger than 5 and when the arrow is yellow”: only left-pointing arrows are presented; c) “press the spacebar when the number is smaller than 5 and when the arrow is yellow”: only right- pointing arrows are presented; d) “press the spacebar when the number is larger than 5 and when the arrow is yellow”: only right - pointing arrows are presented.

The other half of participants performed the same task responding only to blue rather than yellow arrows. Both in the Go-yellow and in the Go-Blue group, the four experimental conditions/blocks of trials were administered during a single experimental session and the order of experimental conditions/blocks of trials was counterbalanced among participants. Each block consisted of 256 trials, 128 with numerical-targets (i.e. 32 trials per Arabic number) and 128 with arrow-targets (64 trials per each one of the two colour conditions). A short break was allowed between blocks. At beginning of the experimental session, participants performed a training session composed of 16 trials.

### 5.3 Results

1,4 % of trials were discarded from the analyses. RTs to numerical targets were entered in an Experimental Condition (Congruent, Incongruent) × Target Magnitude (Smaller, Larger) ANOVA. Neither the Experimental Condition effect [F (1, 19) = 1.53, p = .23, η_p_^2^ = .07] nor the Experimental Condition × Target Magnitude [F (1, 19) = 1.38, p = .25, η_p_^2^= .07; Figure 3A] were significant, showing that also in this task no number-space association (SNAs) was present. The presence of SNAs was also investigated through regression analysis. One-sample t-tests showed that the linear regression slope was not different from zero [t (19) = −1,29, p = .21; average = −2,23; SD = 7.70; Figure 3B]. This confirmed the absence of the SNAs. The counting direction style had no influence on the lack of the SNAs [Group × Experimental Condition interaction: F (1, 16) = .86, p = .36, η_p_^2^ = .04; 10 left to right vs. 10 right to left counting participants].

**Figure 3.**
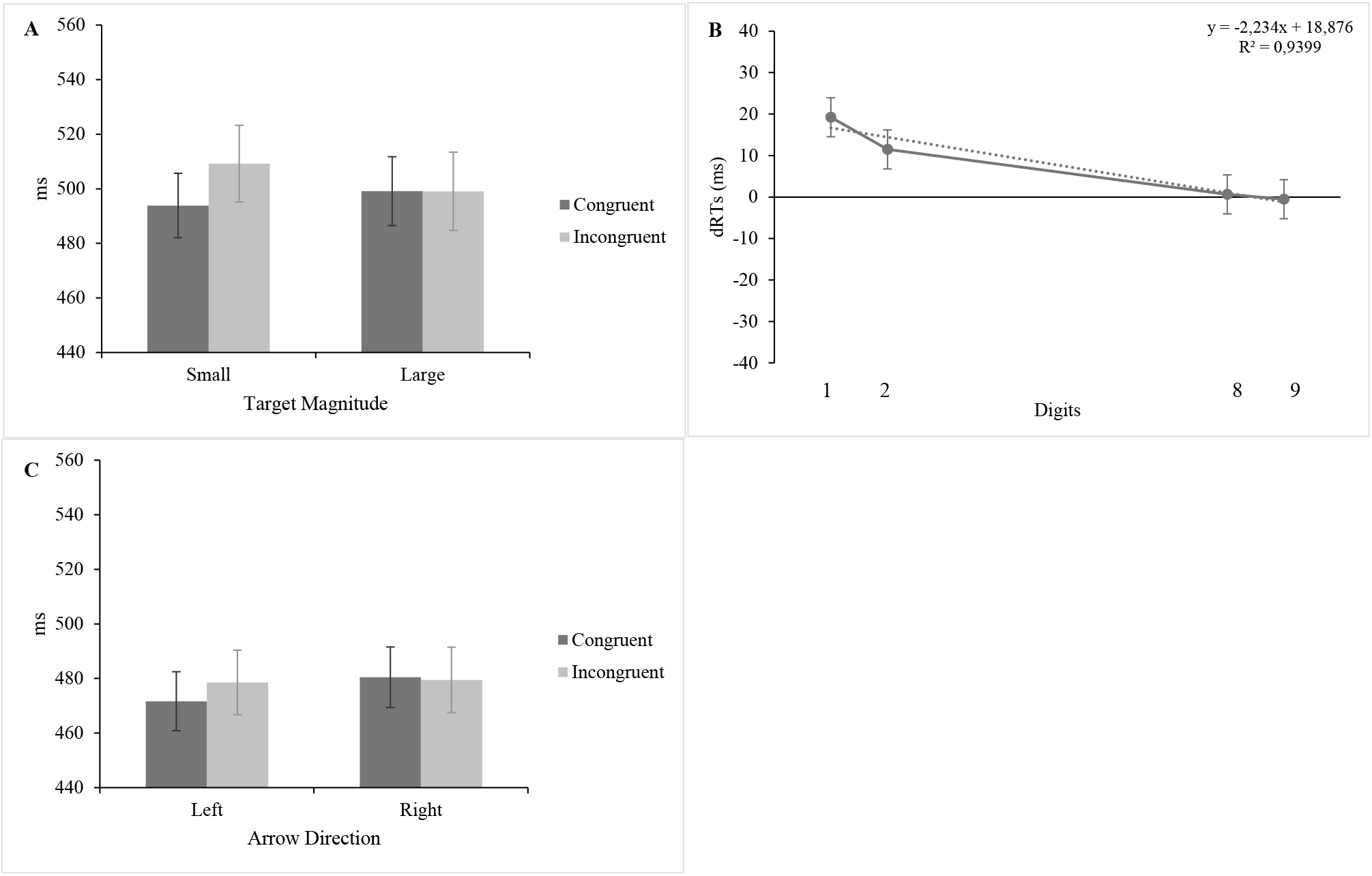
*Experiment 3:* In this experiment participants provided Go responses only to blue or yellow arrow targets. In different block of trials, arrows always pointed in the Left or Right direction. No processing of arrow-direction was required by the task. (**A**) Average RTs (with SE) to Small magnitude (1,2) and Large magnitude (8,9) numerical targets in the Congruent task condition (RTs to Small number magnitudes when in a given block arrows always pointed to the Left and RTs to Large number magnitude when a given block arrows always pointed to the Right) and in the Incongruent condition (RTs to Small number magnitudes when in a given block arrows always pointed to the Right and RTs to Large number magnitude when a given block arrows always pointed to the Left). (**B**) Regression slope describing differential RTs (dRT in ms) to Small magnitude and Large magnitude numerical targets between blocks with Right pointing minus blocks with Left pointing arrows. (**C**) Average RTs (with SE) to Left and Right pointing arrows in the Congruent task condition (RTs to Left arrows when participants must respond only to Small number magnitudes and RTs Right pointing arrows when participants must respond only to Large number magnitudes) and in the Incongruent condition (RTs to Left arrows when participants must respond only to Large number magnitudes and RTs Right pointing arrows when participants must respond only to Small number magnitudes).

Finally, RTs to arrows targets were entered in an Experimental Condition (Congruent, Incongruent) ×Arrow Direction (Left, Right) × Arrow Colour (Yellow, Blue) ANOVA. The Experimental Condition effect was not significant [F (1, 19) = .030, p = .59, η_p_^2^= .01; Congruent: 476 ms; incongruent: 479ms; Figure 3C)]: this showed that, like in experiment 2, attending to numbers lower than 5 did not speed up RTs to arrows pointing left and, vice versa attending to numbers higher than 5 did not speed up RTs to arrows pointing right. Also in this case, the counting direction style had no influence on the lack of the SNAs [Group × Experimental Condition interaction: F (1, 18) = 1.47, p= .24, η_p_^2^= .07].

## 6. Discussion

The set of results reported in the series of experiments summarised the present study shows, quite convincingly, that contrasting left/right spatial codes used in isolation do not elicit a corresponding spatial representation of number magnitudes and, vice versa, contrasting small/larger number magnitude codes used in isolation do not elicit a corresponding activation of left/right contrasting spatial codes.

The results of Experiment 1 provided direct evidence for this conclusion by showing that when coloured arrow-targets that pointed leftward or rightward and numerical-targets that were smaller or larger than 5 were alternated at central fixation: a) providing Go responses to a specific arrow direction (left or right) produced no RTs advantage in the detection of numerical targets that are supposed to be placed in a spatially congruent position on a left-to-right oriented MNL; b) providing Go response to a specific numerical magnitude, smaller or larger than 5, produced no RTs advantage in the detection of arrow-targets that were oriented toward the same position that numbers would putatively occupy on a left-to-right oriented MNL. These results provide a clear response to the first problem that we wished to address in the present study and strongly suggest, in agreement with our original theoretical proposal (Aiello et al., 2012; Fattorini et al., 2015; 2016), that congruency space-number effects like those described in Fischer and Shaki (2017) depend, as in the case of the SNARC effect, on the concomitant use spatial and numerical codes in the instructions that regulate the performance of Go/NoGo responses rather than on an inherent association between space and number magnitude codes.

The results of Experiment 2 confirmed these conclusions by demonstrating that in a task with alternating numerical- and arrow-targets in which no association was present between a specific arrow-colour and a specific arrow-direction, discriminating numerical magnitudes did not speeded up the detection of arrows whose direction was congruent with the putative position occupied by the same numerical magnitudes on a the left-to-right oriented MNL. These results are important, in that they suggest that the number-magnitude/arrow direction congruency effect described by Shaki and Fischer (2018) was likely due to the fact that in their experiment a specific colour was always associated with a specific arrow direction thus inducing, in the mental task set generated by the instruction of attending simultaneously to a specific number-magnitude and a specific arrow-colour, a spurious link between number-magnitude and spatial-codes .

In Experiment 3 we expanded on the results of experiment 2 and tried to verify whether in a task that required the concomitant discrimination of the magnitude of numerical targets and the colour of arrow-targets, the incorporation in the mental task set of the spatial code conveyed by the task irrelevant arrow direction could have been eventually forced when the left or right direction of coloured arrow-targets was maintained constant in a block of trials. The results of this control experiment confirmed the results of experiment 2 and highlighted no implicit activation of spatial codes related to arrow-direction and no arrow direction/number magnitude congruency effects, e.g. when in a block of trials arrow-targets pointed constantly to the left, responses to numbers smaller than 5 were not faster as compared with blocks of trials in which arrow-targets pointed constantly to the right. It is important to note that the evaluation of individual finger counting style direction (Lindemann et al., 2011) demonstrated that, in all experiments, space-number congruency effects were absent both in participants who counted from left-to-right and in those who counted from right-to-left. In addition, all participants showed a left-to-right preference in the sequential visual inspection and pointing of visual items organised in a horizontal row (Fischer and Shaki, 2017).

Taken together, the results of the present study show that a significant and reliable association between space and numbers requires the joint use of contrasting small vs. large magnitude numerical codes and contrasting left vs. right spatial codes. In contrast, space-number associations are not observed when task instructions require the isolated use of contrasting magnitude number codes or the isolated use of contrasting spatial codes. These conclusions diverge from those advanced by Shaki and Fischer (2018) who proposed that the activation of magnitude codes is sufficient in eliciting that of corresponding spatial codes and of a spatially organised MNL, i.e. space-number association. Nonetheless, the results of our experiments 2 and 3 clearly suggest that the conclusion by Shaki and Fischer (2018) is based on the results of an experiment where the activation of spatial codes was spuriously conveyed by the constant association between a specific arrow-colour and a specific arrow-direction.

To summarise, our study provides further evidence in support of our original claim (Aiello et al. 2012; 2013; Fattorini et al., 2015; 2016) that numbers have no inherent spatial organisation and that this organisation is rather elicited when spatial codes are used in the task at hand. At the neural level, the non-overlap between the representation of space and number-magnitude is supported by the finding that the topographical distribution of neuronal population cells that code for different numerosities has no correspondence with that of neuronal populations that code for different sectors of space (Harvey et al., 2013). In the present study, we expand on our original proposal and show that in order to evoke spatially organised MNLs, spatial and numerical codes must be used jointly in the task at hand. These findings have also important methodological implications as they suggest that in order to properly test the spontaneous adoption of a spatially organised mental arrangement of numbers, the use of contrasting left/right or up/down spatial labels or the spatial arrangement of numerical material or response keys along the left/right or up/down reading direction should be accurately avoided in the task set and instructions. A good example of how the spatial arrangement of numerical cues can induce space/number congruency effects that are otherwise absent when cues are not spatially arranged is provided by the finding reported by Longo and Lourenco (2007) and by Rotondaro et al. (2015), respectively. Longo and Laurenco (2007) observed that when healthy participants report the mental subjective midpoint of a numerical intervals by writing it on a horizontal line with the numerical endpoints of the interval printed one to the left and one to the right of the same line, bisection biases toward numbers lower than true midpoints, i.e. supposedly to the left of true numerical midpoints are found and, in the same participants, these biases are correlated to leftward biases that are found in the bisection of horizontal visual lines (spatial pseudo-neglect; Jewell and McCourt, 2000). At variance with this finding, Rotondaro et al. (2015) showed that when the endpoints of number intervals are proposed verbally by the examiner, i.e. with no visual-spatial suggestion biasing their arrangement in mental space, and responded verbally by participants no correlation is longer present between biases in numerical and visual line bisections. These findings are in keeping with the lack of any correlation between numerical and line bisection biases in patients with right brain damage (Zorzi et al., 2002; Doricchi et al., 2005; Doricchi et al., 2009; Aiello et al., 2012; Aiello et al., 2013) and instantiate how in a numerical task the spatial arrangement of numerical cues can induce an equivalent mental spatialization of numbers that is otherwise absent when the same cues are provided to participants without implicit or explicit spatial suggestions.

When discussing the role played by the use of spatial codes in the generation of the SNAs, one should also consider that these codes can be called into action through different cognitive mechanism. As an example, expanding on the role of spatial working memory suggested by the findings in right brain patients reported by Doricchi et al., (2005), van Dijck and co-workers (van Dijck and Fias, 2011; van Dijck et al., 2013; van Dijck et al., 2014) have documented that participants engaged in the active monitoring of the ordinal position of numerical items embedded in short arbitrary sequences of five items presented one by one at central fixation, follow their reading habits and adopt a left-to-right mental organization of these sequences in working memory. This was suggested by the findings of faster RTs to visual targets in the left side of space when these were preceded by the presentation of item belonging to the beginning of the sequence and, vice versa by faster RTs to target in the right side of space when these were preceded by a numerical item positioned at the end of the sequence. Taken together with the findings of our study, these results suggest that SNAs are generated whenever any task-related factor like the use of spatial codes or the memorization of sequences of alphanumerical items, makes easier the performance of numerical judgements through the activation of the visuo-motor and spatial components of acquired reading habits.

In conclusion, as already proposed in Fattorini et al. (2016), the findings and arguments reported in the presented study suggest “*that the widespread conviction, prompted by the reliability of the SNARC effect, that the representation of numbers in the human brain has an inherent directional-spatial component should be profoundly revised*”: additional investigations are suitable to spread further light on the more complex and interesting scenario sketched by the results of our experiments.

## Acknowledgement

We thank Alessandra Martucci for help in data collection. This work was supported by University La Sapienza Grants Grandi Attrezzature Scientifiche (n° C26G119T3J) to FD and Avvio alla Ricerca to MP. FD was also supported by grant Ricerca Corrente 2015 from the Fondazione Santa Lucia IRCCS, Rome.

## References

Aiello, M., Jacquin-Courtois, S., Merola, S., Ottaviani, T., Tomaiuolo, F., Bueti, D., … & Doricchi, F. (2012). No inherent left and right side in human ‘mental number line’: evidence from right brain damage. Brain, aws114. http://dx.doi.org/10.1093/brain/aws114.

Aiello, M., Merola, S., & Doricchi, F. (2013). Small numbers in the right brain: evidence from patients without and with spatial neglect. Cortex, 49(1), 348–351. http://dx.doi.org/10.1016/j.cortex.2012.06.002.

Dehaene, S., Dupoux, E., & Mehler, J. (1990). Is numerical comparison digital? Analogical and symbolic effects in two-digit number comparison. Journal of Experimental Psychology: Human Perception and Performance, 16(3), 626.

Dehaene, S., Bossini, S., & Giraux, P. (1993). The mental representation of parity and number magnitude. Journal of Experimental Psychology: General, 122(3), 371–396. http://dx.doi.org/10.1037/0096-3445.122.3.371.

Doricchi, F., Guariglia, P., Gasparini, M., & Tomaiuolo, F. (2005). Dissociation between physical and mental number line bisection in right hemisphere brain damage. Nature Neuroscience, 8(12), 1663–1665. http://dx.doi.org/10.1038/nn1563.

Doricchi, F., Merola, S., Aiello, M., Guariglia, P., Bruschini, M., Gevers, W., et al. (2009). Spatial orienting biases in the decimal numeral system. Current Biology, 19(8), 682–687. http://dx.doi.org/10.1016/j.cub.2009.02.059.

Fattorini, E., Pinto, M., Rotondaro, F., & Doricchi, F. (2015). Perceiving numbers does not cause automatic shifts of spatial attention. Cortex, 73, 298–316.

Fattorini, E., Pinto, M., Merola, S., D’Onofrio, M., & Doricchi, F. (2016). On the instability and constraints of the interaction between number representation and spatial attention in healthy humans: A concise review of the literature and new experimental evidence. In Progress in brain research (Vol. 227, pp. 223–256). Elsevier.

Fias, W. (1996). The importance of magnitude information in numerical processing: Evidence from the SNARC effect. Mathematical cognition, 2(1), 95–110.

Fischer, M. H., & Shaki, S. (2017). Implicit spatial-numerical associations: Negative numbers and the role of counting direction. Journal of Experimental Psychology: Human Perception and Performance, 43(4), 639.

Gevers, W., Ratinckx, E., De Baene, W., & Fias, W. (2006). Further evidence that the SNARC effect is processed along a dual-route architecture: Evidence from the lateralized readiness potential. Experimental psychology, 53(1), 58–68. doi: http://dx.doi.org/10.1027/1618-3169.53.L58

Gevers, W., Santens, S., Dhooge, E., Chen, Q., Van den Bossche, L., Fias, W., & Verguts, T. (2010). Verbal-spatial and visuospatial coding of number-space interactions. Journal of Experimental Psychology: General, 139(1), 180.

Harvey, B. M., Klein, B. P., Petridou, N., & Dumoulin, S. O. (2013). Topographic representation of numerosity in the human parietal cortex. Science, 341(6150), 1123–1126.

Hubbard, E.M., Piazza, M., Pinel, P., Dehaene, S., 2005. Interactions between number and space in parietal cortex. Nat. Rev. Neurosci. 6 (6), 435–448.

Ishihara, M., Jacquin-Courtois, S., Flory, V., Salemme, R., Imanaka, K., & Rossetti, Y. (2006). Interaction between space and number representations during motor preparation in manual aiming. Neuropsychologia, 44(7), 1009–1016. http://dx.doi.org/10.1016/j.neuropsychologia.2005.11.008.

Ito, Y., & Hatta, T. (2004). Spatial structure of quantitative representation of numbers: Evidence from the SNARC effect. Memory & Cognition, 32(4), 662–673.

Jewell, G., & McCourt, M. E. (2000). Pseudoneglect: a review and meta-analysis of performance factors in line bisection tasks. Neuropsychologia, 38(1), 93–110.

Kadosh, R. C., Lammertyn, J., & Izard, V. (2008). Are numbers special? An overview of chronometric, neuroimaging, developmental and comparative studies of magnitude representation. Progress in neurobiology, 84(2), 132–147.

Keus, I. M., Jenks, K. M., & Schwarz, W. (2005). Psychophysiological evidence that the SNARC effect has its functional locus in a response selection stage. Cognitive Brain Research, 24(1), 48–56. doi:10.1016/j.cogbrainres.2004.12.005

Keus, I. M., & Schwarz, W. (2005). Searching for the functional locus of the SNARC effect: Evidence for a response-related origin. Memory & Cognition, 33(4), 681–695. http://dx.doi.org/10.3758/BF03195335.

Lindemann, O., Alipour, A., & Fischer, M. H. (2011). Finger counting habits in middle eastern and western individuals: an online survey. Journal of Cross-Cultural Psychology, 42(4), 566–578.

Loetscher, T., & Brugger, P. (2009). Random number generation in neglect patients reveals enhanced response stereotypy, but no neglect in number space. Neuropsychologia, 47(1), 276–279.

Loetscher, T., Bockisch, C. J., Nicholls, M. E., & Brugger, P. (2010). Eye position predicts what number you have in mind. Current Biology, 20(6), R264–R265.

Longo, M. R., & Lourenco, S. F. (2007). Spatial attention and the mental number line: Evidence for characteristic biases and compression. Neuropsychologia, 45(7), 1400–1407.

Lorch, R. F., & Myers, J. L. (1990). Regression analyses of repeated measures data in cognitive research. Journal of Experimental Psychology: Learning, Memory, and Cognition, 16(1), 149.

Müller, D., & Schwarz, W. (2007). Exploring the mental number line: evidence from a dual-task paradigm. Psychological Research, 71(5), 598–613. http://dx.doi.org/10.1007/s00426-006-0070-6

Nuerk, H. C., Wood, G., & Willmes, K. (2005). The universal SNARC effect: The association between number magnitude and space is amodal. Experimental psychology, 52(3), 187–194.

Pia, L., Neppi-Modona, M., Cremasco, L., Gindri, P., Dal Monte, O., & Folegatti, A. (2012). Functional independence between numerical and visual space: evidence from right brain damaged patients. Cortex, 48(10), 1351–1358. http://dx.doi.org/10.1016/j.cortex.2012.04.005.

Pinto, M., Fattorini, E., Lasaponara, S., D’Onofrio, M., Fortunato, G., & Doricchi, F. (2018). Visualising numerals: a ERPs study with the Attentional SNARC task. Cortex. https://doi.org/10.1016Zj.cortex.2017.12.015

Proctor, R. W., & Cho, Y. S. (2006). Polarity correspondence: A general principle for performance of speeded binary classification tasks. Psychological bulletin, 132(3), 416.

Rossetti, Y., Jacquin-Courtois, S., Aiello, M., Ishihara, M., brozzolli, C., & Doricchi, F. (2011). Neglect “around the Clock”: dissociating number and spatial neglect in right brain damage. In S. Dehaene, & E. M. Brannon (Eds.), Space, time and number in the brain: Searching for the foundations of mathematical thought (pp. 149–173). Amsterdam: Elsevier. Retrieved from http://dx.doi.org/10.1016/B978-0-12-385948-8.00011-6.

Rotondaro, F., Merola, S., Aiello, M., Pinto, M., & Doricchi, F. (2015). Dissociation between line bisection and mental-number-line bisection in healthy adults. Neuropsychologia, 75, 565–576.

Santens, S., & Gevers, W. (2008). The SNARC effect does not imply a mental number line. Cognition, 108(1), 263–270.

Shaki, S., & Fischer, M. H. (2018). Deconstructing spatial-numerical associations. Cognition, 175, 109–113.

van Dijck, J. P., Gevers, W., Lafosse, C., Doricchi, F., & Fias, W. (2011). Non-spatial neglect for the mental number line. Neuropsychologia, 49(9), 2570–2583. http://dx.doi.org/10.1016/j.neuropsychologia.2011.05.005.

van Dijck, J. P., & Fias, W. (2011). A working memory account for spatial-numerical associations. Cognition, 119(1), 114–119.

van Dijck, J. P., Gevers, W., Lafosse, C., & Fias, W. (2012). The heterogeneous nature of numberespace interactions. Frontiers in Human Neuroscience, 5. http://dx.doi.org/10.3389/fhhum.2011.00182.

van Dijck, J. P., Abrahamse, E. L., Majerus, S., & Fias, W. (2013). Spatial attention interacts with serial-order retrieval from verbal working memory. Psychological Science, 24(9), 1854–1859.

van Dijck, J. P., Abrahamse, E. L., Acar, F., Ketels, B., & Fias, W. (2014). A working memory account of the interaction between numbers and spatial attention. The Quarterly Journal of Experimental Psychology, 67(8), 1500–1513.

Vuilleumier, P., Ortigue, S., & Brugger, P. (2004). The number space and neglect. Cortex, 40(2), 399–410. http://dx.doi.org/10.1126/science.1239052.

Wood, G., Willmes, K., Nuerk, H. C., & Fischer, M. H. (2008). On the cognitive link between space and number: A meta-analysis of the SNARC effect. Psychology Science Quarterly, 50(4), 489.

Zanolie, K., & Pecher, D. (2014). Number-induced shifts in spatial attention: a replication study. Frontiers in Psychology, 5. http://dx.doi.org/10.3389/fpsyg.2014.00987.

Zorzi, M., Priftis, K., & Umiltà, C. (2002). Brain damage: neglect disrupts the mental number line. Nature, 417(6885), 138–139.

Zorzi, M., Bonato, M., Treccani, B., Scalambrin, G., Marenzi, R., & Priftis, K. (2012). Neglect impairs explicit processing of the mental number line. Frontiers in Human Neuroscience, 6. http://dx.doi.org/10.3389/hum.2012.00125.

